# CoVac501, a self-adjuvanting peptide vaccine conjugated with TLR7 agonists, against SARS-CoV-2 induces protective immunity

**DOI:** 10.1101/2021.04.10.439275

**Authors:** Yiru Long, Jianhua Sun, Tingting Liu, Feng Tang, Xinxin Zhang, Qiuping Qin, Yunqiu Miao, Weiliang Zhu, Xiaoyan Pan, Qi An, Mian Qin, Xiankun Tong, Xionghua Peng, Pan Yu, Peng Zhu, Yachun Zhang, Leike Zhang, Gengfu Xiao, Jianping Zuo, Wei Tang, Ji Zhou, Zhijian Xu, Yong Gan, Jin Ren, Wei Huang, Guangyi Jin, Likun Gong

**Affiliations:** State Key Laboratory of Drug Research, Shanghai Institute of Materia Medica, Chinese Academy of Sciences, Shanghai, 201203, China; University of Chinese Academy of Sciences, Beijing, 100049, China; State Key Laboratory of Virology, Wuhan Institute of Virology, Chinese Academy of Sciences, Wuhan, 430071, China; Shanghai King-Cell Biotechnology Co., Ltd., Shanghai, 201500, China; Zhongshan Institute for Drug Discovery, Institutes of Drug Discovery and Development, Chinese Academy of Sciences, Zhongshan, 528400, China; School of Pharmaceutical Sciences, Shenzhen University Health Science Center, Shenzhen University, Shenzhen, 518060, China; International Cancer Center, Nation-Regional Engineering Lab for Synthetic Biology of Medicine, Shenzhen University, Shenzhen, 518060, China; School of Pharmaceutical Science and Technology, Hangzhou Institute for Advanced Study, University of Chinese Academy of Sciences, Hangzhou, 310024, China

## Abstract

Safe, economical and effective vaccines against severe acute respiratory syndrome coronavirus 2 (SARS-CoV-2) are needed to achieve adequate herd immunity and halt the pandemic. We have constructed a novel SARS-CoV-2 vaccine, CoVac501, which is a self-adjuvanting peptide vaccine conjugated with Toll-like receptor 7 (TLR7) agonists. The vaccine contains two immunodominant peptides screened from receptor-binding domain (RBD) and is fully chemically synthesized. And the vaccine has optimized nanoemulsion formulation, outstanding stability and safety. In non-human primates (NHPs), CoVac501 elicited high and persistent titers of RBD-specific and protective neutralizing antibodies (NAbs), which were also effective to RBD mutations. CoVac501 was found to elicit the increase of memory T cells, antigen-specific CD8^+^ T cell responses and Th1-biased CD4^+^ T cell immune responses in NHPs. More importantly, the sera from the immunized NHPs can prevent infection of live SARS-CoV-2 in vitro.

**One-Sentence Summary:** A novel SARS-CoV-2 vaccine we developed, CoVac501, which is a fully chemically synthesized and self-adjuvanting peptides conjugated with TLR7 agonists, can induce high-efficient humoral and cellular immune responses against SARS-CoV-2.

## Main Text

Safe, quickly designable, protective and economical vaccines against severe acute respiratory syndrome coronavirus 2 (SARS-CoV-2) are required to rapidly respond to viral mutations, reduce the economic burden on public health and halt coronavirus disease 2019 (COVID-19) epidemics (*1–3*). Epitope-specific peptide vaccines can be rapidly designed against viral proteins, readily respond to mutations in pathogens and completely chemically synthesized on a large scale with low production cost (*4, 5*).

However, the immunogenicity of peptides is relatively low (*6*). Coupling innate immune agonists to form self-adjuvanting vaccine is a promising way to improve the immunogenicity of peptide vaccines and to elicit specific antigen presentation. Toll-like receptor 7 (TLR7) is a natural immune pattern recognition receptor localized on the endo/lysosomes membrane (*7*). TLR7 agonists are being developed in combination with various vaccines in our and others’ studies to facilitate the antigen presentation, activate innate immunity, promote antibody affinity maturation and maintain long-lasting immune memory (*8–11*).

We have developed a novel SARS-CoV-2 vaccine, named CoVac501, which is a fully chemically synthesized and self-adjuvanting peptide vaccine conjugated with TLR7 agonists. CoVac501 can induce high-level, durable and protective neutralizing antibodies (NAbs) against SARS-CoV-2 and RBD mutations and induce specific T cell immune responses in non-human primates (NHPs).

### Prediction and screening of peptide candidates

Spike (S) glycoprotein mediates SARS-CoV-2 entry into host cells through the interaction between receptor-binding domain (RBD) and angiotensin-converting enzyme 2 (ACE2)(*12, 13*). Several studies have shown that the RBD is an immunodominant region in the S protein and NAbs in the sera of convalescent patients are mostly against the RBD (*14–16*). Screening of neutralizing antibody epitopes from RBD for immunization holds promise for inducing specific and protective humoral immune responses.

Based on computer-aided vaccine design technology, we integrated the sequence and structural features of RBD to predict and screen several immunodominant peptide candidates (Fig. 1A).

**Fig. 1.**
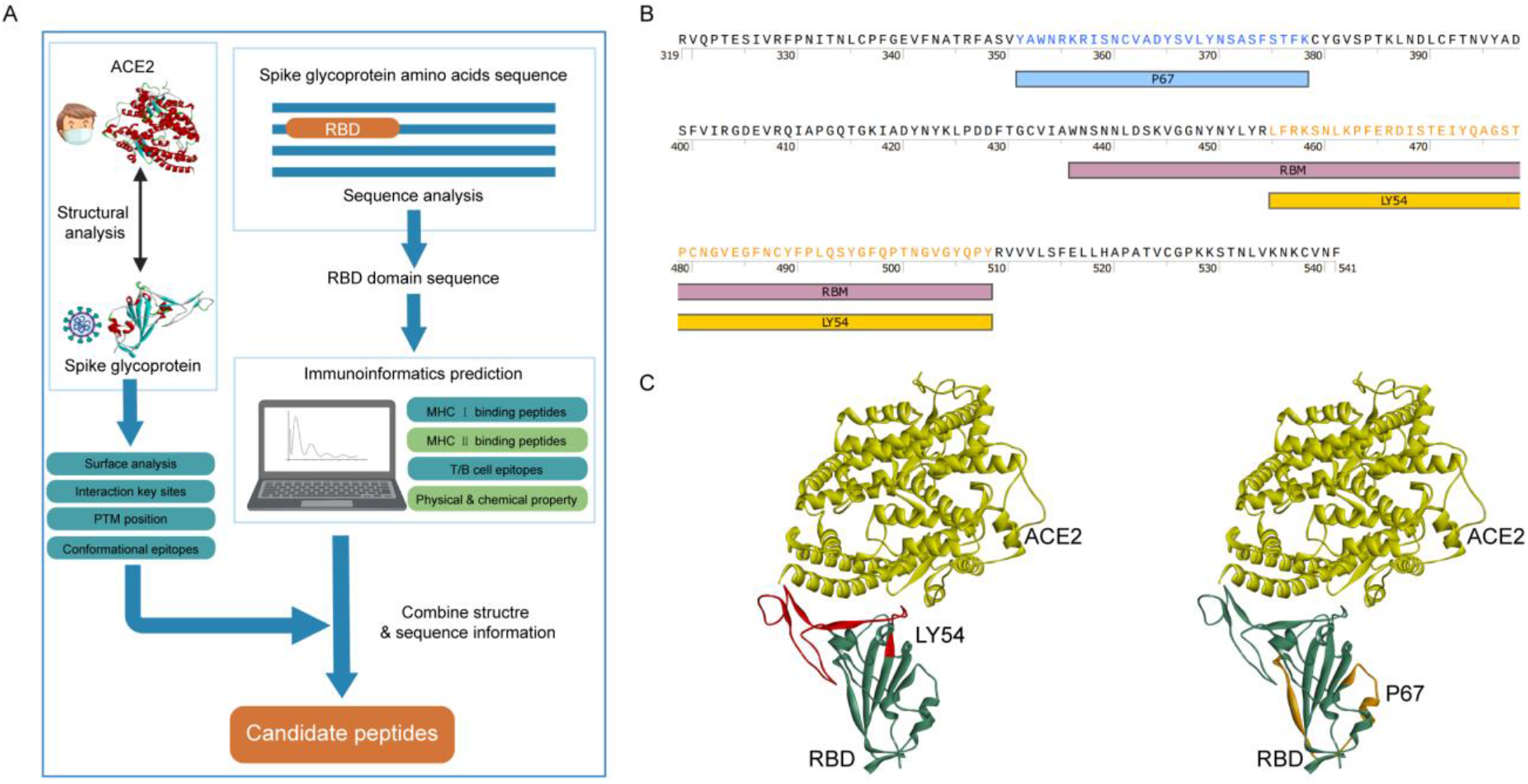
Identification of immunodominant candidate peptides. (**A**) The flowchart illustrates the prediction and screening process of the RBD immunodominant peptides. The candidate peptides were determined by combining the sequence and structural features of the RBD. The flowchart was created with BioRender.com. (**B**) The graph shows that the amino acid sequences of candidate peptides LY54 and P67. RBM, receptor binding motif. (**C**) The graph shows that the spatial localization of candidate peptides LY54 and P67 in the RBD. The crystal structure of SARS-CoV-2 RBD bound with ACE2 was from Protein Data Bank (PDB; code 6M0J). The red color shows the location of LY54 and the orange color shows the location of P67.

Based on NetMHCIIpan-4.0 (*17*), we predicted human leukocyte antigen (HLA) class II binding peptides for major HLA class II types (fig. S1, A and B). The RBD contains three immunodominant regions: residues around 345-380, 450-470 and 485-515. Consistently, these regions also contain potential HLA class I epitopes and B-cell linear epitopes predicted by NetMHCpan-4.1 (*17*) and BepiPred-2.0 (*18*)(fig. S1, C and D). Considering the physicochemical properties, we selected LY54 (residues 455-508) and P67 (residues 351-378) as candidate peptides (Fig. 1B and fig. S1E). And LY54 is positioned at the interaction interface between RBD and ACE2 (Fig. 1C). By peptide solid-phase synthesis technique, we prepared LY54 and P67 with high purity.

### Preparation and identification of peptides conjugated with TLR7 agonists

By conjugating TLR7 agonists, peptides can form self-adjuvanting vaccines, which will allow antigen-presenting cells (APCs) to simultaneously uptake peptide antigens and TLR7 agonist adjuvants to induce a potent antigen-specific immune response (*19*). We have developed a TLR7 small molecule agonist, named SZU-101, which has been applied in our previous studies for the construction of self-adjuvanting vaccines (*8, 9*). To conjugate LY54 and P67 with SZU-101, the N-Hydroxysuccinimide (NHS) esters group and the maleimide (Mal) group were covalently conjugated to the carboxyl group of SZU-101, respectively (Fig. 2A). LY54-101 were prepared by coupling SZU-101-NHS with the leucine (residue 1) and lysine (residues 4 and 8) of LY54 (Fig. 2A). SZU-101-Mal was attached to the cysteine (residue 12) of P67 via a sulfhydryl group to form P67-101 (Fig. 2A). LY54-101 and P67-101 were purified and characterized by high performance liquid chromatography (HPLC) and high-resolution mass spectra (HRMS) to achieve more than 95% purity (fig. S2 to S7).

**Fig. 2.**
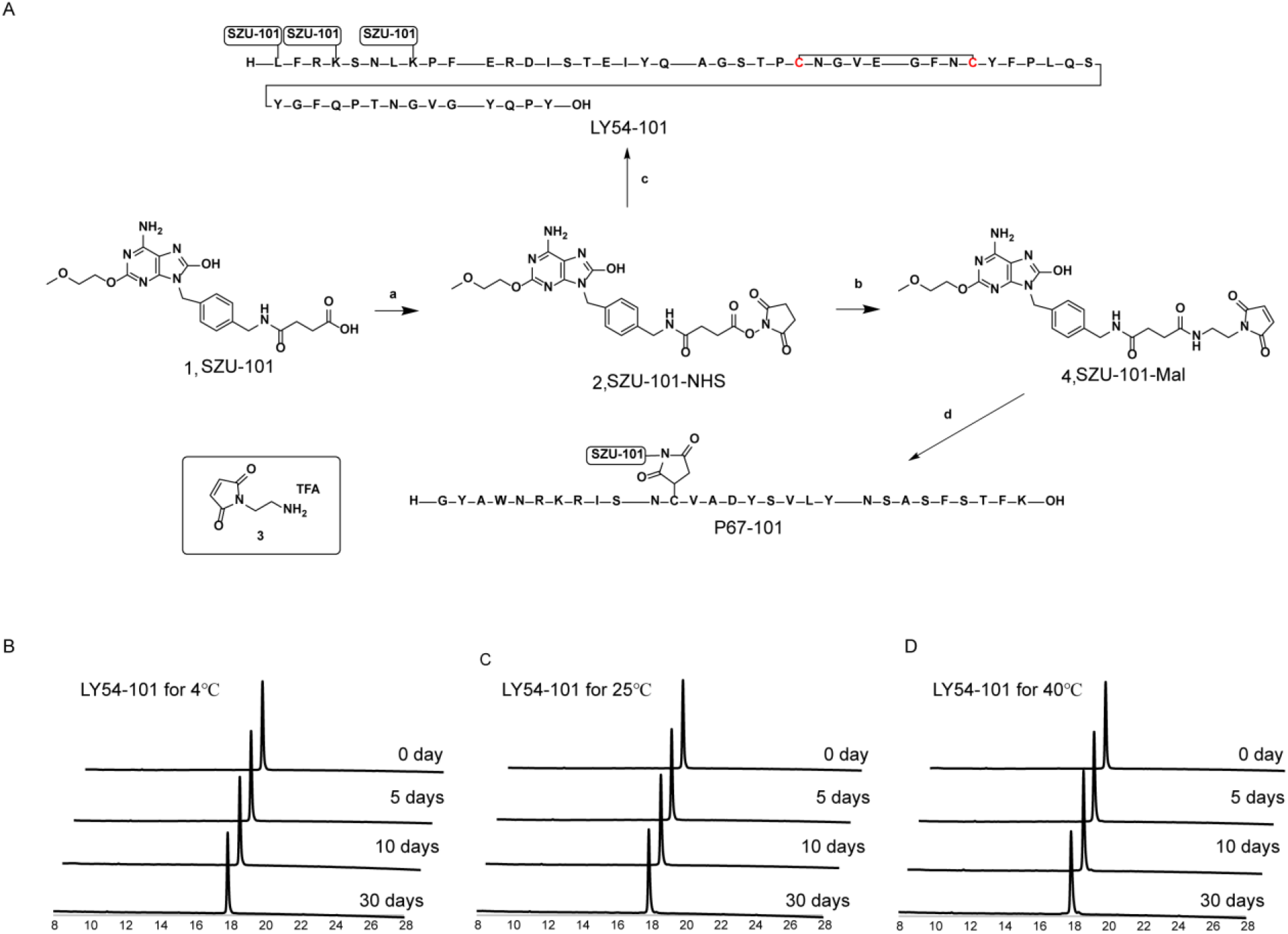
Full chemical synthesis and thermal stability analysis of self-adjuvanting peptide vaccines. (**A**) Synthesis of SZU-101 derivatives and SZU-101-peptide conjugations. The synthesis of LY54 and P67 was performed by solid phase peptide synthesis technique. Reaction conditions: a, EDCI/NHS, DMSO, 15°C, overnight; b, 3, EDCI/NHS, DMSO, 15°C, overnight; c, LY54, NaHCO_3_, DMF/H_2_O, room temperature (rt), 2h; d, P67, NaHCO_3_, DMF/H_2_O, rt, 1h. (**B** to **D**) HPLC profile of stability analysis of LY54-101 for 5 days, 10 days and 30 days at 4°C (B), 25°C (C) and 40°C (D). The horizontal axis represents the retention of time (min) and the vertical axis represents the absorbance (mAU).

Considering that the thermal stability of vaccines can greatly affect the shipping and storage conditions, we tested the thermal stability of LY54-101 and P67-101. LY54-101 and P67-101 were placed at 4°C, 25°C and 40°C for 5 days, 10 days and 30 days, respectively. HPLC identification results showed that no significant changes were found in the purity of LY54-101 and P67-101 at 4°C, 25°C and even 40°C for 30 days, suggesting that LY54-101 and P67-101 has outstanding thermal stability (Fig. 2, B to D).

In summary, we prepared high quality peptides conjugated with TLR7 agonists (LY54-101 and P67-101) with prominent thermal stability through full chemical synthesis approach.

### Immunogenicity of LY54-101 and P67-101

To preliminary evaluate the immunogenicity of LY54-101 and P67-101, C57BL/6 mice were injected with LY54-101 (50 μg) and P67-101 (50 μg) mixed with Titermax adjuvant for three doses at days 0, 7 and 14. The sera obtained 20 days after the third vaccination showed a strongly specific antibody response against RBD in the mice, with IgG titer up to 1: 1000000 (Fig. 3A). Notably, the sera obtained 56 days after the third vaccination still showed a RBD specific IgG titer up to 1: 400000 (Fig. 3B). Further, the immunogenicity of LY54-101 and P67-101 was accessed in NHP models. Cynomolgus monkeys were injected with LY54-101 (1mg) and P67-101 (1mg) mixed with Titermax adjuvant for two doses at days 0 and 14 (Fig. 3C). The sera obtained 20 days post the second vaccination showed a strongly specific antibody response against RBD in the cynomolgus monkeys, with IgG titer up to 1: 24300 (Fig. 3D) and a NAbs titer greater than 1:160 (Fig. 3E). Notably, the sera obtained 91 days after the second vaccination still showed a RBD specific IgG titer up to 1: 24300 (Fig. 3F) and a NAbs titer greater than 1:320 (Fig. 3G), suggesting that the self-adjuvanting peptide vaccine induced a potent and long-lasting humoral immune response.

**Fig. 3.**
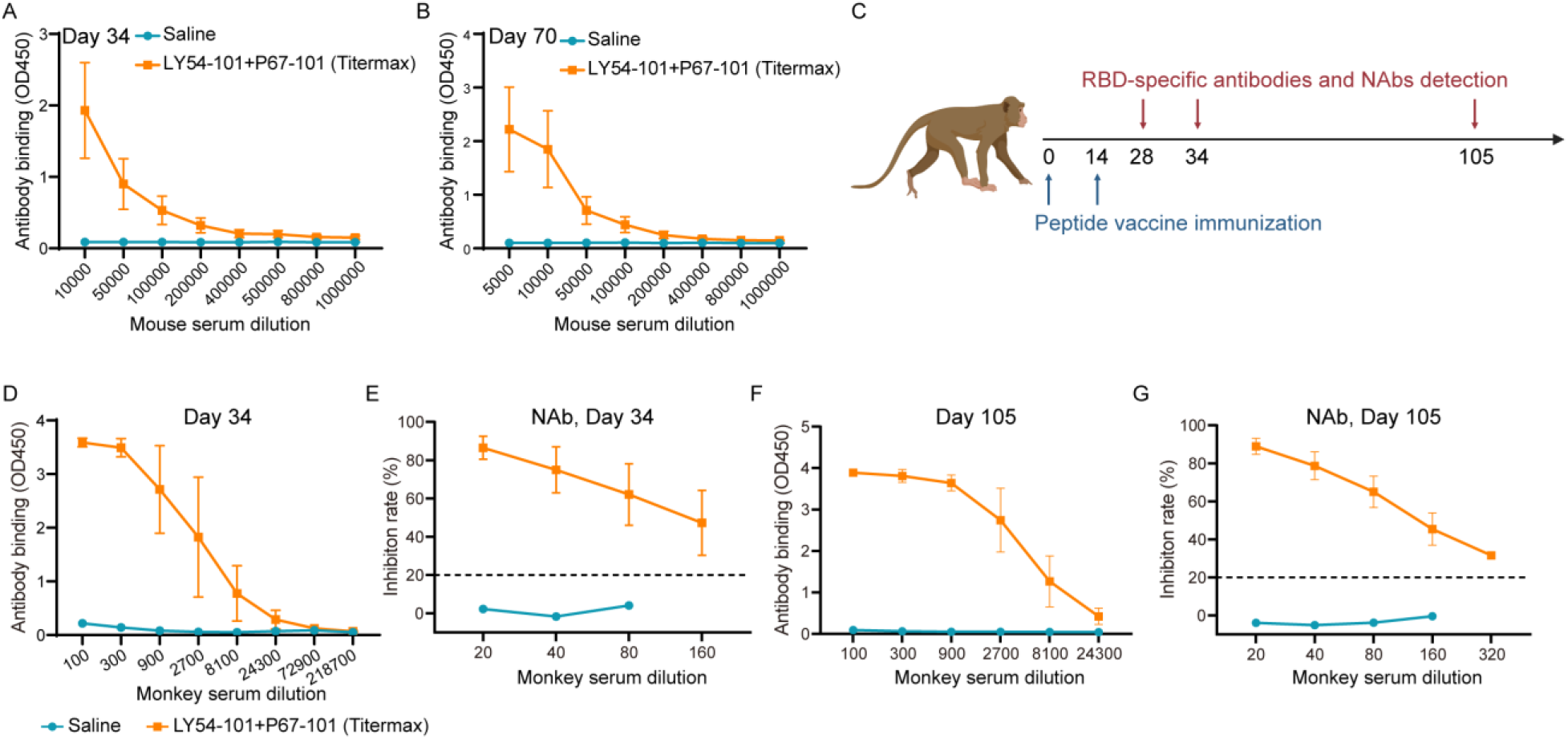
Immunogenicity of LY54-101 and P67-101. (**A** and **B**) The C57BL/6 mice (n = 5) were immunized with LY54-101 and P67-101 in the presence of Titermax for three intramuscular injections on days 0, 7 and 14. Sera were collected from the mice 34 days (A) and 70 days (B) after the first dose of vaccine and the levels of IgG against the recombinant RBD protein were tested for different serum dilutions using ELISA. The data are presented as the mean ± SEM. (**C**) Schematic diagram of immunization, sample collection and detection in cynomolgus monkeys. (**D** and **E**) The cynomolgus monkeys (n = 2) were immunized with LY54-101 and P67-101 in the presence of Titermax for two intramuscular injections on days 0 and 14. Sera were collected from the monkeys 34 days after the first dose of vaccine and the levels of RBD-specific antibody (D) and NAbs (E) were tested for different serum dilutions using ELISA. A 20% inhibition rate is the cut-point for a positive titer of NAbs. The data are presented as the mean ± SEM. (**F** and **G**) Sera were collected from the monkeys 105 days after the first dose of vaccine and the levels of RBD-specific antibody (F) and NAbs (G) were tested for different serum dilutions using ELISA. The data are presented as the mean ± SEM.

### Vaccine formulation optimization and in vivo delivery

Adjuvant formulations suitable for our self-adjuvanting peptide vaccine and available for clinical application need to be developed to replace Titermax. Nanoemulsions formulations can protect vaccine antigens, achieve slow release of the antigen and promote antigenic load of lymph nodes, and have been widely used in vaccine development, such as AS03 (*20*). We developed two nanoemulsion formulations (F1 and F2) and compared their adjuvant effects with AS03 and Titermax for the in vivo delivery of our self-adjuvanting peptide vaccines.

To study the retention of vaccine formulation at injection sites, several Cy5 labeled LY54-101 formulations were intramuscularly injected into the upper inner thigh of rats, and the fluorescence signal in inner thighs over 8 h was investigated using a live animal fluorescence imaging system (Fig. 4A). Free LY54-101 showed little fluorescence signal 4 h post injection, while LY54-101 nanoemulsions showed enhanced retention at injection sites (Fig. 4B). F2 nanoemulsion displayed slightly decreased fluorescence signal compared to AS03 nanoemulsion, with more than 50% of fluorescence intensity remained at the injection sites (Fig. 4B).

**Fig. 4.**
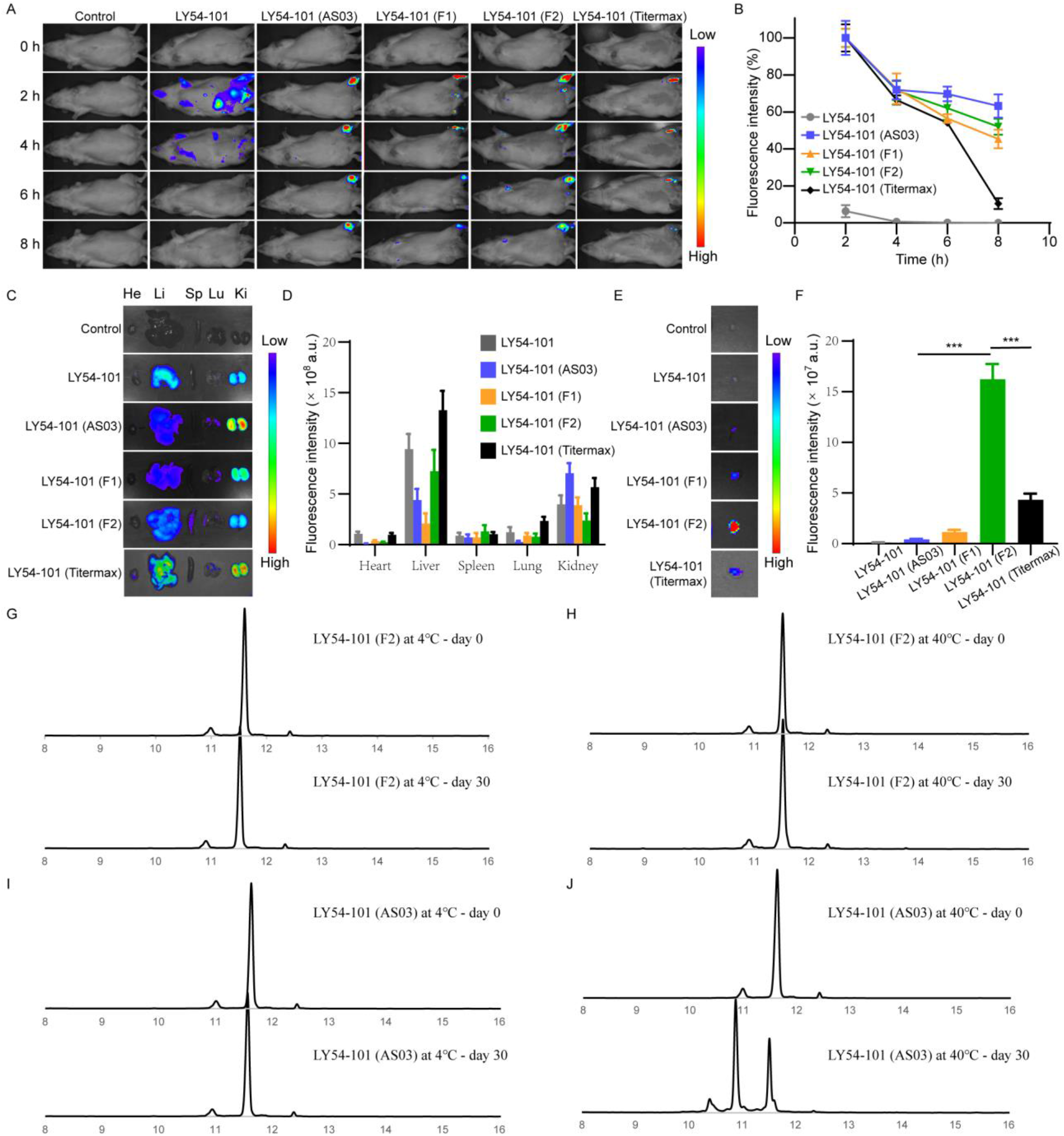
Bioditribution and retention of peptide vaccine nanoemulsions in vivo. (**A**) Real-time in vivo fluorescence images of rats after intramuscular injection of Cy5 labeled LY54-101 nanoemulsions. (**B**) Monitoring of the kinetics of LY54-101 nanoemulsion retention at injection sites within 8 h. (n = 3; mean ± SD). (**C**) Fluorescence images of organs excised at 8 h postinjection. (**D**) Biodistribution of Cy5 labeled LY54-101 nanoemulsions in main organs excised from rats at 8 h postinjection. (n = 3; mean ± SD). (**E**) Fluorescence images of lymph nodes excised at 8 h postinjection. (**F**) Fluorescence intensity of Cy5 labeled LY54-101 nanoemulsions in lymph nodes excised from rats at 8 h postinjection. (n = 3; mean ± SD), *** P<0.001 as determined by one-way ANOVA with multiple comparison tests. (**G** to **J**) HPLC profile of stability analysis of F2 nanoemulsion formulation (G and H) and AS03 nanoemulsion formulation (I and J) for LY54-101 at 4°C or 40°C for 30 days. The horizontal axis represents the retention of time (min) and the vertical axis represents the absorbance (mAU).

Major organs were excised from rats 8 h post injection to study the biodistribution of different LY54-101 formulations. Free LY54-101 and LY54-101 nanoemulsions showed strong fluorescence signals both in liver and kidney, indicating that LY54-101 was mainly metabolized through liver and kidney (Fig. 4, C and D). We next focused on the recruitment of different LY54-101 formulations in lymph nodes. Little free LY54-101 was recruited in lymph node, while AS03, F1 and F2 nanoemulsions enhanced the recruitment of LY54-101 in lymph node (Fig. 4E). Notably, F2 nanoemulsion displayed the highest fluorencence intensity in lymph node among all vaccine formulations including the Titermax (Fig. 4F). We speculated that F2 nanoemulsions for LY54-101 would obtain better immune efficacy than AS03 nanoemulsions.

In addition, we tested the thermal stability of vaccine formulations. F2 nanoemulsion formulation and AS03 nanoemulsion formulation for LY54-101 were placed at 4°C or 40°C for 30 days, respectively. HPLC identification results showed that no significant changes were found in the purity of LY54-101 in both F2 nanoemulsion formulation and AS03 nanoemulsion formulation at 4°C for 30 days (Fig. 4, G and I). Notably, LY54-101 in the AS03 nanoemulsion formulation was degraded after one month at 40°C, while LY54-101 in the F2 nanoemulsion formulation remained stable, revealing that the F2 nanoemulsion formulation of LY54-101 has outstanding stability (Fig. 4, H and J).

### Humoral immune responses in CoVac501 vaccinated Cynomolgus macaques

To determine the optimized formulation composition, study the role of the TLR7 agonist in self-adjuvanting peptide vaccines and determine the ratios optimized for LY54-101 and P67-101, we vaccinated cynomolgus monkeys with peptide vaccines in the following groups: saline, LY54-101 (F2), LY54-101 (AS03), LY54-101+P67-101 (1:1, F2), LY54-101+P67-101 (2:1, F2) and LY54+P67 (1:1, F2) (Fig. 5A). Animals received peptide vaccines via the intramuscular route for three doses at days 0, 14 and 28, while were found to have RBD-specific binding antibodies determined by an enzyme-linked immunosorbent assay (ELISA) and NAbs determined by both a competitive ELISA and a live virus neutralization assay.

**Fig. 5.**
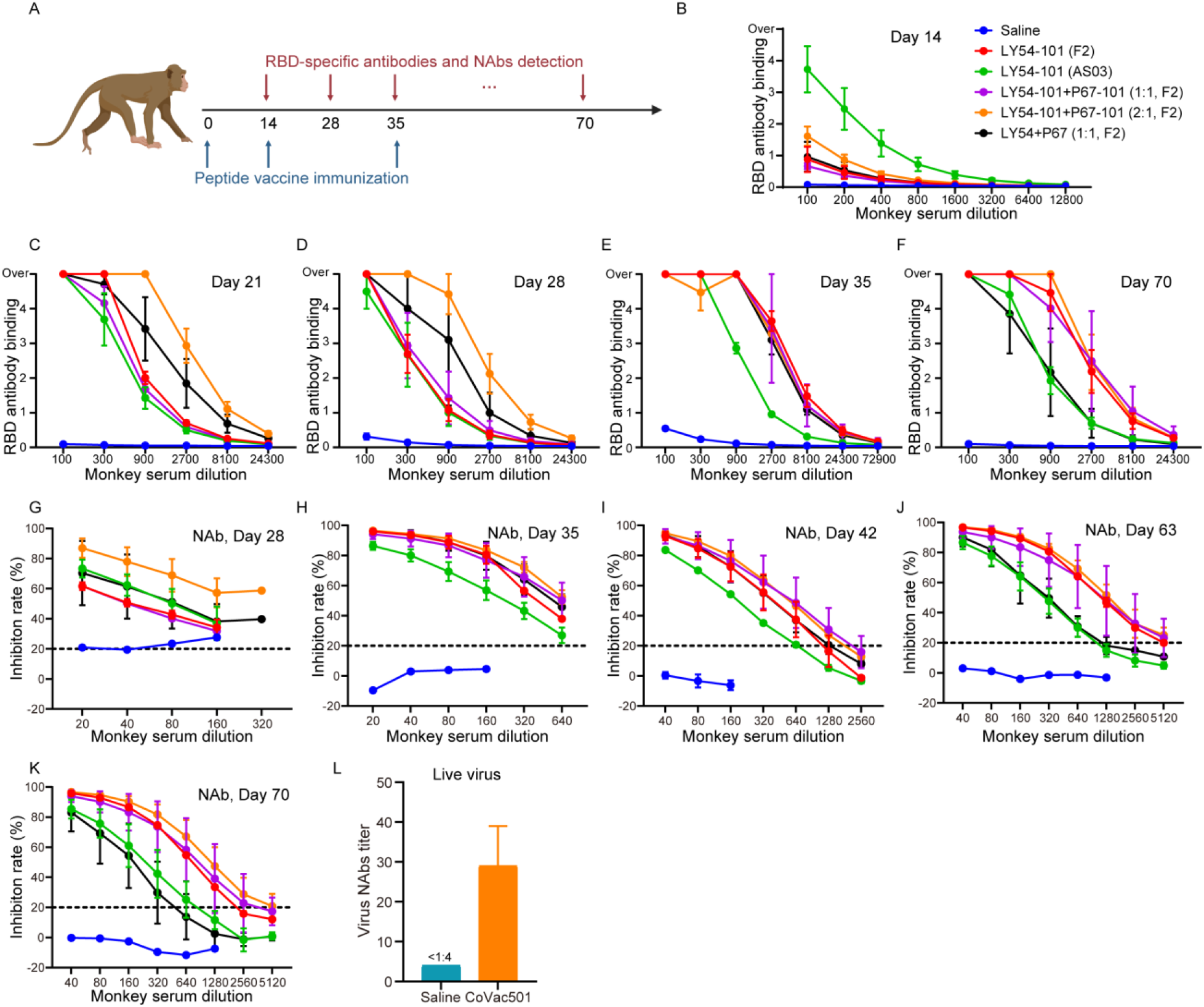
CoVac501 induced protective humoral immune responses in Cynomolgus macaques. (**A**) Schematic diagram of immunization, sample collection and detection in cynomolgus monkeys. (**B** to **F**) The cynomolgus monkeys (n = 2) were immunized with peptide vaccines in the following groups: saline, LY54-101 (F2), LY54-101 (AS03), LY54-101+P67-101 (1:1, F2), LY54-101+P67-101 (2:1, F2) and LY54+P67 (1:1, F2) for three doses at days 0, 14 and 28. Sera were collected from the monkeys 14 (B), 21 (C), 28 (D), 35 (E) and 70 (F) days after the first dose of vaccine and the levels of RBD-specific antibody were tested for different serum dilutions using ELISA. The data are presented as the mean ± SEM. (**G** to **K**) Sera were collected from the monkeys 28 (G), 35 (H), 42 (I), 63 (J) and 70 (K) days after the first dose of vaccine and NAbs were tested for different serum dilutions using ELISA. The data are presented as the mean ± SEM. (**L**) The neutralization of sera from cynomolgus monkeys 56 days after the first dose of vaccine for live SARS-CoV-2 in vitro was tested by microdose cytopathogenic efficiency (CPE) assay. The data are presented as the mean ± SEM.

Sera obtained on day 14 after the first dose of peptide vaccines already showed specific IgG responses to the RBD, especially for the LY54-101 (AS03) group (Fig. 5B). The titer of RBD-specific binding antibodies gradually increased with two booster immunizations from day 14 to day 35 and peaked between day 35 and day 42, while a decreasing trend was observed from day 42 to day 70 (Fig. 5, C to F and fig. S8). By comparing the changes in binding antibody levels between the groups, we determined the composition of the vaccine formulation. First, comparing the binding antibody titers in the LY54-101 (F2) and LY54-101 (AS03) groups, we found that AS03 nanoemulsion could induce humoral immune responses more rapidly, but the later antibody levels were much lower than those induced by F2 nanoemulsion, suggesting that F2 nanoemulsion is more suitable for our peptide vaccines. Second, P67-101 further increased the production of specific antibodies to RBD, thus the optimal immunization ratio of LY54-101 and P67-101 was 2:1. Third, by conjugating with the TLR7 agonist SZU-101, the peptide vaccines induced a stronger and longer-lasting humoral immune response. Therefore, we identified LY54-101+P67-101 (2:1, F2) as the final self-adjuvanting peptide vaccine formulation and named it CoVac501.

Notably, the neutralizing antibody titer continued to increase and was maintained at a stable level from day 63 to day 70, which was eventually up to 1:5120 (Fig. 5, G to K). Further, we tested the neutralization of sera from vaccinated cynomolgus monkeys for live SARS-CoV-2 in vitro by microdose cytopathogenic efficiency (CPE) assay. Sera from CoVac501 vaccinated cynomolgus monkeys had a neutralization titer up to 1:39 at 28 days after the third immunization (Fig. 5L). These results indicated that the CoVac501 can induce a high level, protective and durable humoral immune response.

### Humoral immunity induced by CoVac501 against RBD mutations

Mutations in amino acid residues of the RBD may affect the effectiveness of existing vaccines and neutralizing antibodies. We therefore tested changes in antibody titers of sera from CoVac501 vaccinated cynomolgus monkeys against five mutant RBD proteins (N439K, Y453F, S477N, E484K and N501Y) (Fig. 6, A to D). Sera were obtained on days 35, 56 and 70 after the first dose of CoVac501. The results showed that RBD-specific antibody levels was not affected by mutations of N439K, Y453F, S477N and N501Y of RBD. E484K mutation reduced the RBD-specific antibody levels by threefold, but still retained enough RBD-binding antibodies. Thus, the binding ability of RBD-specific antibodies induced by CoVac501 was not significantly affected by mutations at multiple loci on the RBD.

**Fig. 6.**
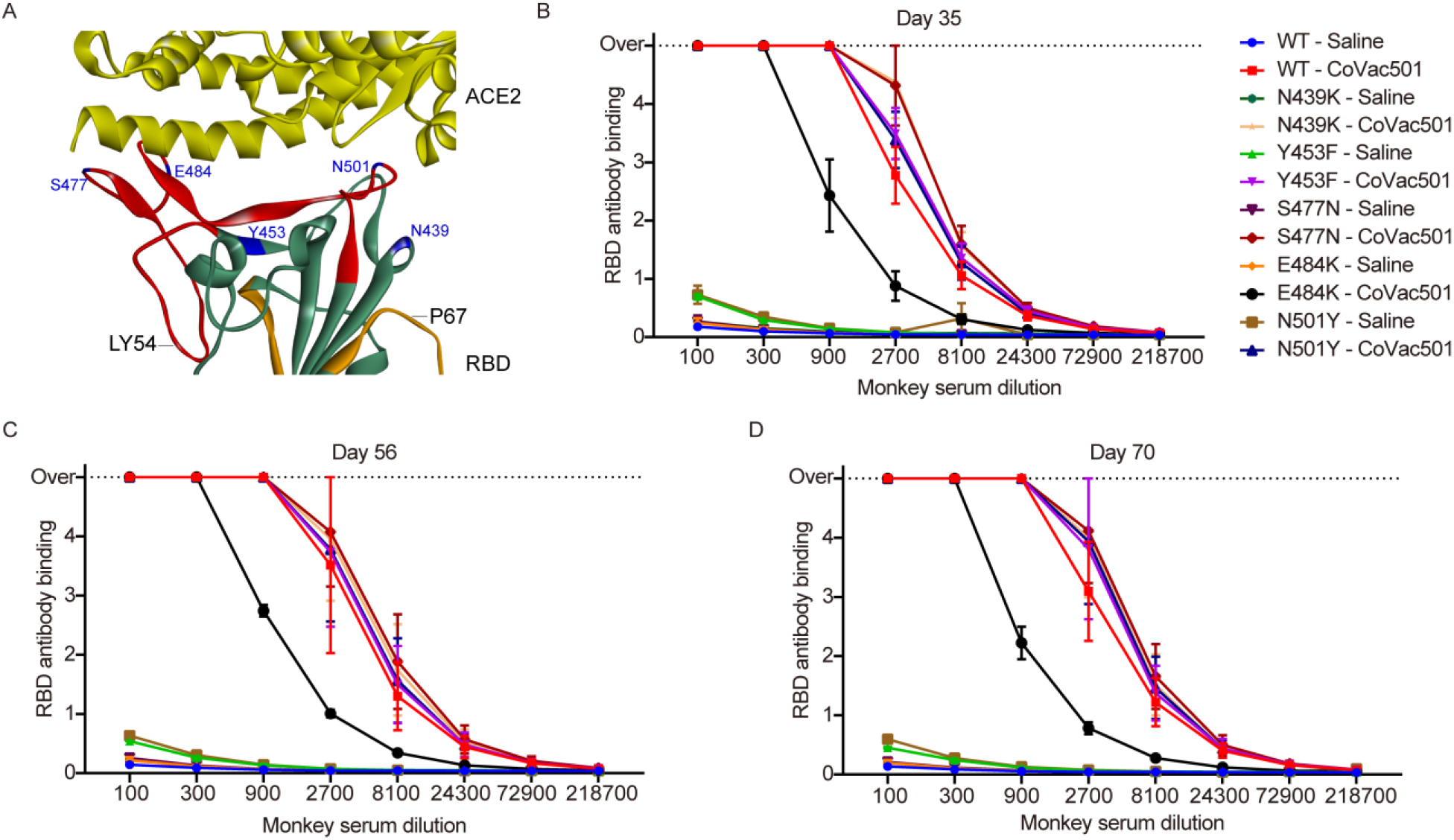
CoVac501 induced humoral immune responses against RBD mutations. (**A**) Positioning of N439K, Y453F, S477N, E484K and N501Y in RBD. (**B** to **D**). The cynomolgus monkeys (n = 2) were immunized with vehicle or CoVac501 for three doses at days 0, 14 and 28. Sera were collected from the monkeys 35 (B), 56 (C) and 70 (D) days after the first dose of vaccine and the levels of RBD mutations (N439K, Y453F, S477N, E484K and N501Y) binding antibodies were tested for different serum dilutions using ELISA.

### T cell immune responses in CoVac501 vaccinated Cynomolgus macaques

We also observed cellular immune response in CoVac501 vaccinated monkeys. T cells of vaccinated monkeys (day 28 and day 42) were quantified by flow cytometry. CoVac501 vaccinated monkeys had fewer naive T cells (CD28^+^, CCR7^+^ and CD45RA^+^), more effector T cells (CD28^-^, CCR7^-^ and CD45RA^+^) and more memory T cells (CD28^+^, CCR7^-^and CD45RA^-^), suggesting that CoVac501 vaccination would promote T-cell activation and T-cell immune memory in cynomolgus monkeys (Fig. 7, A and B). In addition, PBMCs of vaccinated monkeys were stimulated in vitro with either LY54 peptides or RBD. Intracellular cytokine staining assays showed that CoVac501 induced an RBD or LY54-specific CD8^+^T cells responses (Fig. 7, C to D). Meanwhile, CoVac501 induced an increase in IFN-γ^+^, TNF-α^+^ and IL-2^+^ CD4^+^ T cells without affecting the levels of IL-4^+^, IL-6^+^ and IL-10^+^ CD4^+^ T cells, indicating that CoVac501 induced Th1-biased immune responses in vaccinated monkeys (Fig. 7, E to F).

**Fig. 7.**
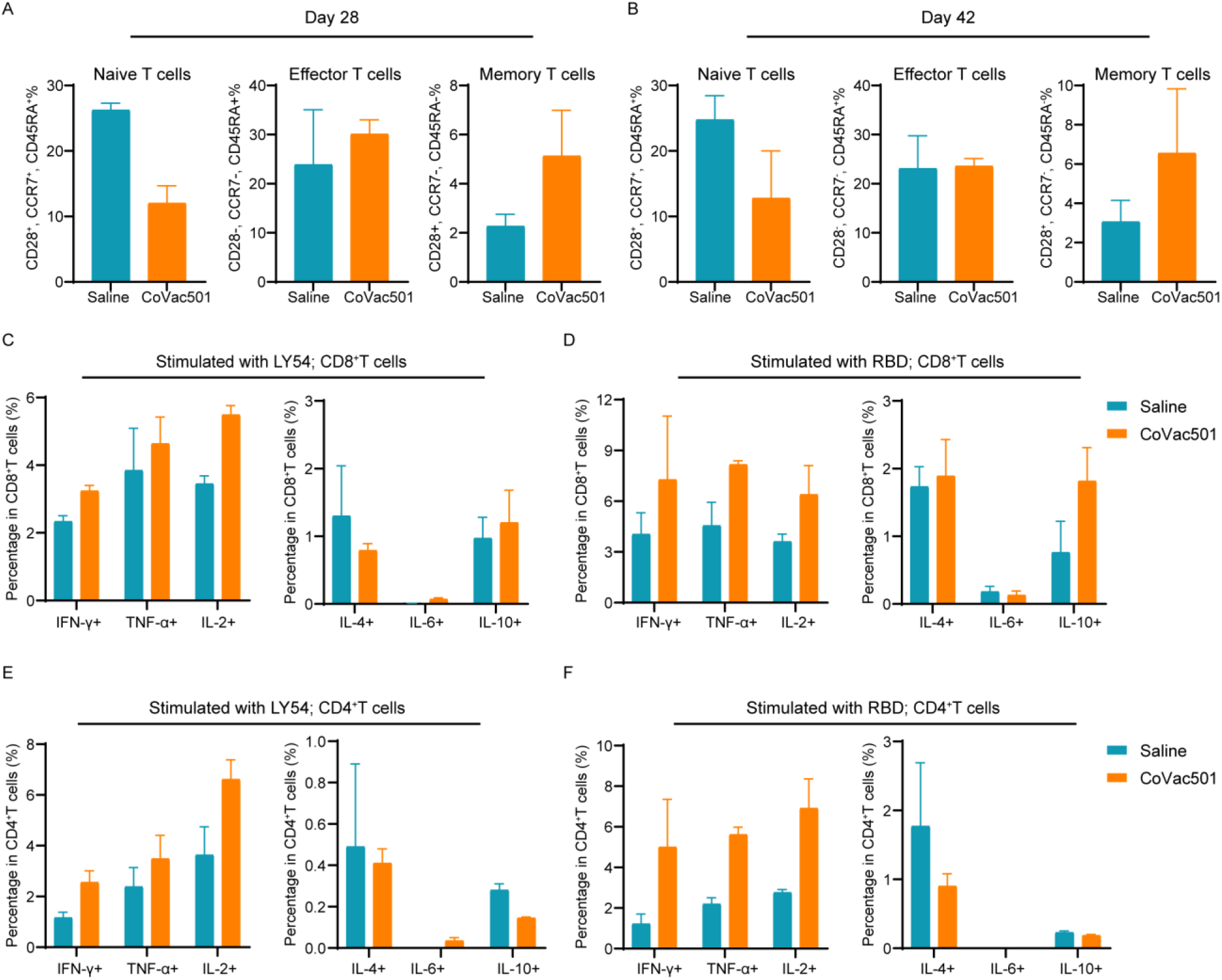
CoVac501 induced a Th1-baised responses and T-cell immune memory in Cynomolgus macaques. (**A** and **B**) PBMCs were collected from the monkeys 28 (A) and 42 (B) days after the first dose of vaccine and naive T cells (CD28^+^, CCR7^+^ and CD45RA^+^), effector T cells (CD28^-^, CCR7^-^ and CD45RA^+^) and memory T cells (CD28^+^, CCR7^-^ and CD45RA^-^) were tested by flow cytometry. (**C** to **E**) PBMCs were collected from the monkeys 70 days after the first dose of vaccine and IFN-γ^+^, TNF-α^+^, IL-2^+^, IL-4^+^, IL-6^+^ and IL-10^+^ intracellular cytokine staining assays by flow cytometry for CD8^+^ (C and D) and CD4^+^ T (E and F) cells in response LY54 (C and E) or RBD (D and F) after 8h antigen stimulation. The data are presented as the mean ± SEM.

## Discussion

The self-adjuvanting peptide vaccine conjugated with TLR7 agonists, represented by CoVac501 we developed, is a novel form of vaccine that has the potential to be applied for COVID-19 or other infectious diseases and can serve as a rapid response strategic reserve for future outbreaks.

CoVac501 is an epitope-based peptide vaccine. Based on computer-aided vaccine design technology, peptide vaccines can be rapidly designed for antigenic proteins. CoVac501 contains two RBD antigen peptides, namely LY54 and P67, which can elicit a neutralizing humoral immune response. Several studies of SARS-CoV-2 NAbs development and spike protein immunogenicity analysis have shown that the regions where LY54 and P67 are located contain the most critical immunodominant and neutralizing epitopes in RBD, suggesting the rationality of our computer-based peptide vaccine design (*14, 21–25*) Moreover, antigenic peptides can be easily and economically produced on a large scale by full chemical synthesis, allowing peptide vaccines to respond rapidly to pandemics and virus mutations. In addition, the main components of CoVac501 have outstanding thermal stability, which makes the vaccine easier to store and transport.

TLR7 agonists have been used as adjuvants in vaccine development (*26*). In vaccine formulations, TLR7 agonists can activate innate immunity, facilitate the antigen presentation process, promote antibody affinity maturation and maintain long-lasting immune memory (*7, 11, 27, 28*). CoVac501 induced a sustained increase in NAbs titers and maintained a durable humoral immune response, which may be attributed to TLR7 agonists for promoting antibody affinity maturation and immune memory formation. Interestingly, the genome of SARS-CoV-2 is linear single-stranded positive-sense RNA (ssRNA), which is recognized by TLR7 in innate immune cells (*12, 29, 30*). Studies have shown that TLR7 expression levels and activity are lower in males, the elderly or people with underlying diseases such as obesity, which are SARS-CoV-2 susceptible populations (*31, 32*). This suggests a potential association between susceptibility to SARS-CoV-2 and TLR7 expression levels and activity. Adequate activation of TLR7 may also provide protection to susceptible individuals (*33*).

The RBD of SARS-CoV-2 has been undergoing mutations. Particularly, 501Y.V1 variant (B.1.1.7) and 501Y.V2 (B.1.351) have been widely spread and have demonstrated the potential for immune escape against existing vaccines and neutralizing antibody drugs (*34–37*). There is an urgent need to develop vaccines against the SARS-CoV-2 mutant strain. Serum antibodies induced by CoVac501 was effective to RBD with mutations in residues N439K, Y453F, S477N and N501Y, suggesting that the 501Y.V1 variant may not show immune escape against CoVac501. Although the mutation of E484K reduced the titer of RBD-specific antibodies induced by CoVac501 by threefold, on the one hand CoVac501 retained enough antibodies against the E484K RBD and on the other hand peptide vaccines can be rapidly designed to cope with the mutation of E484K.

Our present study has some limitations that will be addressed in further research. First, the specific role and mechanism of action of TLR7 agonist SZU-101 in the immune effect of CoVac501 remains to be investigated, especially how TLR7 agonists promote antibody affinity maturation and immune memory formation in humoral immunity in vivo. Second, the components and involving mechanism related to lymph node aggregation and better in vivo immune effect of the peptide vaccine caused by F2 nanoemulsion rather than AS03 nanoemulsion also remains to be elucidated.

In conclusion, we have developed a novel SARS-CoV-2 vaccine CoVac501, which is a self-adjuvanting peptide vaccine conjugated with TLR7 agonists. CoVac501 is a fully chemically synthesized vaccine with outstanding thermal stability and optimized nanoemulsion formulation. CoVac501 can induce high level, protective and long-lasting neutralizing humoral immune responses and Th1-biased T-cell immune responses in NHPs. Besides, CoVac501 was resistant to multiple mutations of amino acid residues in RBDs. CoVac501 holds great potential for further clinical development and application.

## Acknowledgments

We thank the staff of the Center for Drug Safety Evaluation and Research (CDSER), Shanghai Institute of Materia Medica (SIMM) for the safety evaluation and help with animals husbandry and experiments. We thank Conjugenix Pharma-tech Co., Ltd. (Shenzhen, China) for assisting molecule synthesis.

## Author contributions

L.G., G.J., W.H., Y.G. and J.R. designed the study. Y.L. performed the prediction and screening of peptide candidates and prepared the manuscript. L.G., G.J. and Q.Q. identified candidate peptides. T.L. and M.Q. performed the immunogenicity assay of peptide vaccines. J.S. performed the clinical care and vaccination of the animals. J.Z. performed the synthesis of SZU-101. F.T. performed the synthesis, characterization and stability analysis of LY54-101 and P67-101. Y.M. and X.Z. performed the preparation and optimization of vaccine nanoemulsion formulations. J.S., Y.L. and X.P. performed the T cell immune response assay. X.P., G.F., L.Z., X.T., J.Z., W.T., Y.Z. and Q.A. performed the neutralization assays for pseudoviruses and live viruses. W.Z. and Z.X. assisted the analysis of RBD structural features.

## Data and materials availability

All data are available in the manuscript or the supplementary materials. Correspondence and requests for materials should be addressed to L.G.

## Supplementary Materials

Materials and Methods

Figs. S1 to S8

References

## Supplementary Materials for

### Materials and Methods

#### Animals and study design

Animal husbandry was performed as previous work (*1*). All animal studies were conducted in an Association for Assessment and Accreditation of Laboratory Animal Care International (AAALAC) accredited Good Laboratory Practice (GLP) facility. Standard primate diet (purchased from Beijing Keao xieli Feed) was provided daily in amounts appropriate for the ages and sizes of the animals. Reverse osmosis water was made available ad libitum to each animal via an automatic watering device. The animals were provided additional supplements as a form of environmental enrichment and were given various cage-enrichment devices. The animals were maintained on a 12-hr light/12-hr dark cycle in rooms maintained at 20°C to 26°C with a relative humidity of 40% to 70%.

In the immunogenicity evaluation of peptide vaccines in mice, female C57BL/6 mice (n = 5) were injected intramuscularly with LY54-101 (50 μg) and P67-101 (50 μg) mixed with Titermax adjuvant or saline for three doses at days 0, 7 and 14. Sera were collected at regular intervals for antibody detection.

In the first immunogenicity evaluation of peptide vaccines in monkeys, cynomolgus monkeys (n = 2) were injected intramuscularly with LY54-101 (1mg) and P67-101 (1mg) mixed with Titermax adjuvant or saline for two doses at days 0 and 14. Sera were collected at regular intervals for antibody detection.

In the second immunogenicity evaluation of peptide vaccines in monkeys, we vaccinated cynomolgus monkeys with peptide vaccines in the following groups: saline, LY54-101 (2mg, F2), LY54-101 (2mg, AS03), LY54-101+P67-101 (1:1, 2mg+2mg, F2), LY54-101+P67-101 (2:1, 2mg+1mg, F2) and LY54+P67 (1:1, 2mg+2mg, F2) for three doses at days 0,14 and 28. Sera and PBMCs were collected at regular intervals for antibody detection and immune response assays.

#### Prediction of immunodominant peptide epitopes

The amino acid sequence of the Spike protein of SARS-CoV-2 was obtained from the Uniprot database (P0DTC2). The sequence of the RBD was extracted from residues 319 to 541 of the Spike protein. The human leukocyte antigen (HLA) class II binding peptides of RBD for major HLA class II types (DRB1*07:01, DRB1*03:01, DRB1*15:01, DRB1*11:01, DRB1*01:01, DRB1*13:02, DRB1*13:01, DRB1*04:01, DRB1*11:04, DRB1*04:04, DRB1*15:02, DRB1*09:01, DRB1*14:01, DRB1*01:02, DRB1*15:03, DRB1*12:01, DRB1*04:05, DRB1*10:01, DRB1*04:07 and DRB1*04:03) were predicted by NetMHCIIpan-4.0 (*2*). The HLA class I binding peptides of RBD for major HLA class I types were predicted by NetMHCpan-4.1 (*2*). B-cell linear epitopes of RBD were predicted by BepiPred-2.0 (*3*). Crystal structure of SARS-CoV-2 RBD bound with ACE2 was from Protein Data Bank (PDB; code 6M0J).

#### Synthesis of SZU-101 derivatives and SZU-101-peptide conjugations

##### Reagents and instrument

SZU-101 was synthesized as previously reported work (*4*). EDCI, NHS and 1-(2-Aminoethyl)-1H-pyrrole-2,5-dione 2,2,2-trifluoroacetate were purchased from Bidepharm (Shanghai, China). DMF, DMSO and NaHCO_3_ were purchased from Sinopharm (Shanghai, China). Trifluoroacetic acid (TFA) and acetonitrile (ACN) were purchased from J&K (Shanghai, China). Peptides LY54 and P67 were synthesized by JYMedtech (Shenzhen, China). All chemicals and solvents were used without further purification unless indicated. ^1^H and ^13^C NMR spectra were recorded on a 400 MHz, 500 MHz or 600 MHz instrument. HPLC was performed on a Thermo U3000 instrument equipped with a C18 column (Thermo, Acclaim^™^ 120, 5 μm, 4.6 × 250 mm). Compounds were purified by C18 preparative column (Waters, OBD, 5 μm, 10 × 250 mm) with proper eluent gradient. High-resolution mass spectra (HRMS) were performed on Waters Xevo G2-XS QTof spectrometer.

##### Synthesis of SZU-101-NHS

SZU-101 (60 mg, 0.135 mmol, 1.0 eq), EDCI (64.76 mg, 0.34 mmol, 2.5 eq) and NHS (105.65 mg, 0.92 mmol, 6.75 eq) were dissolved in 1.5 mL DMSO and the solution was stirred at 15°C. The reaction was monitored by analytical HPLC. After 14 h, the reaction mixture was poured into water and the product SZU-101-NHS was precipitated as a white solid from the solution. After 3 repeated wash with water, SZU-101-NHS was obtained as a white powder after lyophilization and was used without further purifications.

##### Synthesis of SZU-101-Mal

SZU-101 (60 mg, 0.135 mmol, 1.0 eq), EDCI (64.76 mg, 0.34 mmol, 2.5 eq) and NHS (105.65 mg, 0.92 mmol, 6.75 eq) were dissolved in 1.5 mL DMSO. The solution was stirred at 15°C and was monitored by analytical HPLC. After 14 h, compound 3 (69 mg, 0.27 mmol, 2.0 eq) and 338 μL of 1 M NaHCO_3_ were added into the reaction mixture. The reaction was stirred at rt for another 1 h before purified by preparative C18 column and the product was obtained as a white powder after lyophilization.

##### Synthesis of LY54-101

Peptide LY54 (816 mg, 0.133 mmol, 1.0 eq) was dissolved in a solution of DMF (20 mL) and water (5 mL). SZU-101-NHS (360.45 mg, 5.0 eq.) was added to above solution and a final concentration of 20-25 mM NaHCO_3_ was added. The reaction mixture was incubated at room temperature for 2 h and was monitored by reverse HPLC. The product was purified by preparative C18 column and was obtained as a white powder after lyophilization. Yield, 820 mg, 83.33%. HRMS, calculated for C_339_H_468_N_88_O_98_S_2_, [M+5H]^5+^ 1481.6835, found, 1481.6805; [M+6H]^6+^ 1234.9042, found, 1234.8986; [M+7H]^7+^ 1058.6333, found, 1058.6230.

##### Synthesis of P67-101

Peptide P67 (160 mg, 1.0 eq) was dissolved in DMF (6.0 mL) and water (4.0 mL), and purified SZU-101-Mal (35.2 mg, 1.3 eq) was added to above solution. A final concentration of 30-35 mM NaHCO_3_ was added to the reaction mixture. The reaction mixture was incubated at room temperature for 1 h before purified by preparative C18 column. The product was obtained as a white powder after lyophilization. Yield 69.52%. HRMS, calculated for C_177_H_253_N_49_O_51_S, [M+4H]^4+^ 979.2186, found 979.2466; [M+5H]^5+^ 783.5764, found 783.5913; [M+6H]^6+^ 653.1483, found 653.1462.

#### Thermal stability analysis

Certain amount (0.3-0.8 mg) of conjugate was weighted into bottle. All bottles containing samples were separated into 3 groups and were stored at 4 °C, 25 °C or 40 °C respectively. One bottle of each group was taken out at day 0, day 5, day 10, day 30 for stability analysis. To the bottles to be analyzed was added DMSO to dissolve the samples, giving a concentration of 0.5 mg/mL. For HPLC analysis, 10 μL of aliquots was subjected and the column was eluted by a linear gradient of 2-90% acetonitrile in 30 min at 1.0 mL/min, 40 °C.

#### Preparation and Characterization of vaccine nanoemulsions

Peptide vaccines were dissolved in squalene (2.1%) and alpha-tocopherol (2.4%) to form a homogeneous oil phase, which accounted for 4.5% by volume in nanoemulasion. Tween-80 as an emulsifier was dissolved in oil phase and mixed uniformly. The PBS buffer as water phase was mixed with the oil phase using the Scientz-IId Ultrasonic Cell Disruptor (Ningbo scientz biotechnology Co., China) to form AS03 nanoemulsions with PDC vaccine concentration of 2 mg/mL. To enhance solubility of peptide vaccines, the amount of squalene increased to 2.5% to prepare F1 nanoemulsions. All other materials and procedures were the same as described in the preparation of AS03 nanoemulsions. To improve retention of peptide vaccines at the injection site, polylactic acid (PLA) block copolymer was added to water phase to prepare F2 emulsion on the basis of F1 nanoemulsions prescription. The particle size of three nanoemulsions was determined by a Malvern Zetasizer Nano ZS analyzer (Worcestershire, UK).

#### Bioditribution and retention of vaccine nanoemulsions in vivo

To evaluate the retention of peptide vaccines nanoemulsions at injection site, Cy5 labeled peptide vaccines nanoemulsions were intramuscularly injected into the upper inner thigh of rats after hair removal. At 0, 2, 4, 6 and 8 h postinjection, the rats were imaged by the IVIS Spectrum System (Caliper Corp. Waltham, Massachusetts, USA). After 8 h of injection, the main organs (hearts, livers, spleens, lungs, and kidneys) and lymph nodes were collected for imaging to study the biodistribution of nanoemulsions in tissues. The fluorescence intensity of nanoemulsions at injection sites and various tissues was analyzed quantitatively with a region of interest (ROI) tool.

#### RBD binding antibody assay

RBD binding antibodies were measured by standard ELISA methods. 96-well ELISA plates (Nunc) were coated with RBD-His (1 μg/mL) in PBS overnight at 4°C. After being washed three times with PBST buffer (PBS with 0.05% Tween-20), 96-well plates were blocked with 200 μL of 1% BSA solution for one hours at 37°C with shaking set at 650 rpm. After being washed, 96-well plates were added with 100 μL of serial dilutions of sera and incubated for one hour at 37°C with shaking set at 650 rpm. After being washed, 96-well plates were incubated with 100 μL of Protein A-horseradish peroxidase (HRP, GenScript, 1:5000) for one hour at 37°C with shaking set at 650 rpm. After being washed, 96-well plates were incubated with 100 μL of tetramethyl benzidine (TMB) substrate solution (KPL) at 37°C with shaking set at 650 rpm for 20 min. The reaction was terminated by adding 2 M of sulfuric acid solution, and the absorbance value was measured at 450 nm with an automatic microplate reader SpectraMax (Molecular Devices).

#### Neutralizing antibody assay

NAbs for blocking RBD/ACE2 interaction were detected by the SARS-CoV-2 sVNT Kit (GenScript). Briefly, the samples and controls are pre-incubated with the HRP-RBD to allow the binding of the circulating neutralization antibodies to HRP-RBD. The mixture is then added to the capture plate which is pre-coated with the hACE2 protein. The unbound HRP-RBD as well as any HRP-RBD bound to non-neutralizing antibody will be captured on the plate, while the circulating neutralization antibodies/HRP-RBD complexes remain in the supernatant and get removed during washing. After washing steps, TMB solution is added, making the color blue. By adding Stop Solution, the reaction is quenched and the color turns yellow. This final solution can be read at 450 nm in a microtiter plate reader. The absorbance of the sample is inversely dependent on the titer of the anti-SARS-CoV-2 neutralizing antibodies.

#### Live virus neutralization assay

Cytopathogenic efficiency (CPE) assay was performed as previous work (*5*). The SARS-CoV-2 virus was seeded and propagated in VERO E6 cells. VERO E6 cells were acquired from the American Type Culture Collection (ATCC) and cultured in Dulbecco’s Modified Eagle Medium (DMEM; Meilunbio) supplemented with 10% heat-inactivated fetal bovine serum (FBS; Life Technologies) and 1% penicillin/streptomycin (Invitrogen). Serum samples were heat-inactivated for 30 minutes at 56°C; two-fold serial dilutions, starting from 1:4, were then mixed with an equal volume of viral solution containing 100 of 50% tissue culture infective dose (TCID50) for SARS-CoV-2. The serum-virus mixture was incubated for 1 hour at 37°C in a 5% CO_2_ humidified atmosphere. After incubation, 100 μL of the mixture at each dilution was added in duplicate to a cell plate containing a semi-confluent VERO E6 monolayer. The plates were incubated for 4 days at 37°C. After 4 days of incubation, cytopathic effect (CPE) of each well was recorded under microscopes. The highest serum dilution that protected more than the 50% of cells from CPE was taken as the neutralization titer.

#### T cell immune immunophenotyping

PBMCs were isolated from cynomolgus monkeys blood by lymphocyte separation solution (Dakewei) and added to 96-well plates (1×10^6^ cells/mL) for culture. Samples were firstly blocked with 4% FBS and anti-CD16/CD32 (Fc γ RIII/Fc γ RII, 2.4G2, BD Biosciences), then incubated with surface marker antibodies for 25 minutes at 4°C and then permeabilized with BD Cytofix/Cytoperm buffer before intracellular labeling antibodies were added for 30 minutes at 4°C. Flow cytometry analysis was performed using ACEA NovoCyte. Cellular events were first gated by forward and side scatter characteristics and then by characteristic fluorescence. Data processing was done through NovoExpress software. Antibody staining of cells for flow cytometry analysis was performed following the antibody manufacturer’s recommendations. The antibodies used in T cell immunophenotyping include FITC Mouse Anti-Human CD3_ε_ Clone SP34 (BD Biosciences), APC Mouse Anti-Human CD28 Clone CD28.2 (BD Biosciences), PE-Cy7 Mouse Anti-Human CD45RA Clone 5H9 (BD Biosciences) and Brilliant Violet 650 anti-human CD197 (CCR7) Antibody (Biolegend).

#### T cell immune responses assay

PBMCs were isolated from cynomolgus monkeys blood by lymphocyte separation solution and added to 96-well plates (1×10^6^ cells/mL) for culture. PBMCs and 40 μg/mL of LY54 or RBD were co-incubated for 8 hours and cytokine secretion was simultaneously blocked with BFA (BD Biosciences). Samples were firstly blocked with 4% FBS and anti-CD16/CD32 (Fc γ RIII/Fc γ RII, 2.4G2), then incubated with surface marker antibodies for 25 minutes at 4°C and then permeabilized with BD Cytofix/Cytoperm buffer before intracellular labeling antibodies were added for 30 minutes at 4°C. Flow cytometry analysis was performed using ACEA NovoCyte. Cellular events were first gated by forward and side scatter characteristics and then by characteristic fluorescence. Data processing was done through NovoExpress software. Antibody staining of cells for flow cytometry analysis was performed following the antibody manufacturer’s recommendations. The antibodies used in intracellular cytokine staining include Hu/NHP CD3 Epsilon FITC (BD Biosciences), Hu/NHP CD4 BV421 L200 (BD Biosciences), Hu CD8 APC RPA-T8 (BD Biosciences), Hu IFN-Gma PE-Cy7 4S.B3 (BD Biosciences), Hu TNF BV650 Mab11 (BD Biosciences), Hu IL-2 BV605 MQ1-17H12 (BD Biosciences), Hu/NHP IL-6 PE MQ2-6A3 (BD Biosciences), APC Mouse Anti-Human IL-4 (BD Biosciences) and anti-human IL-10 BV421 (BD Biosciences).

---

**Fig. S1.**
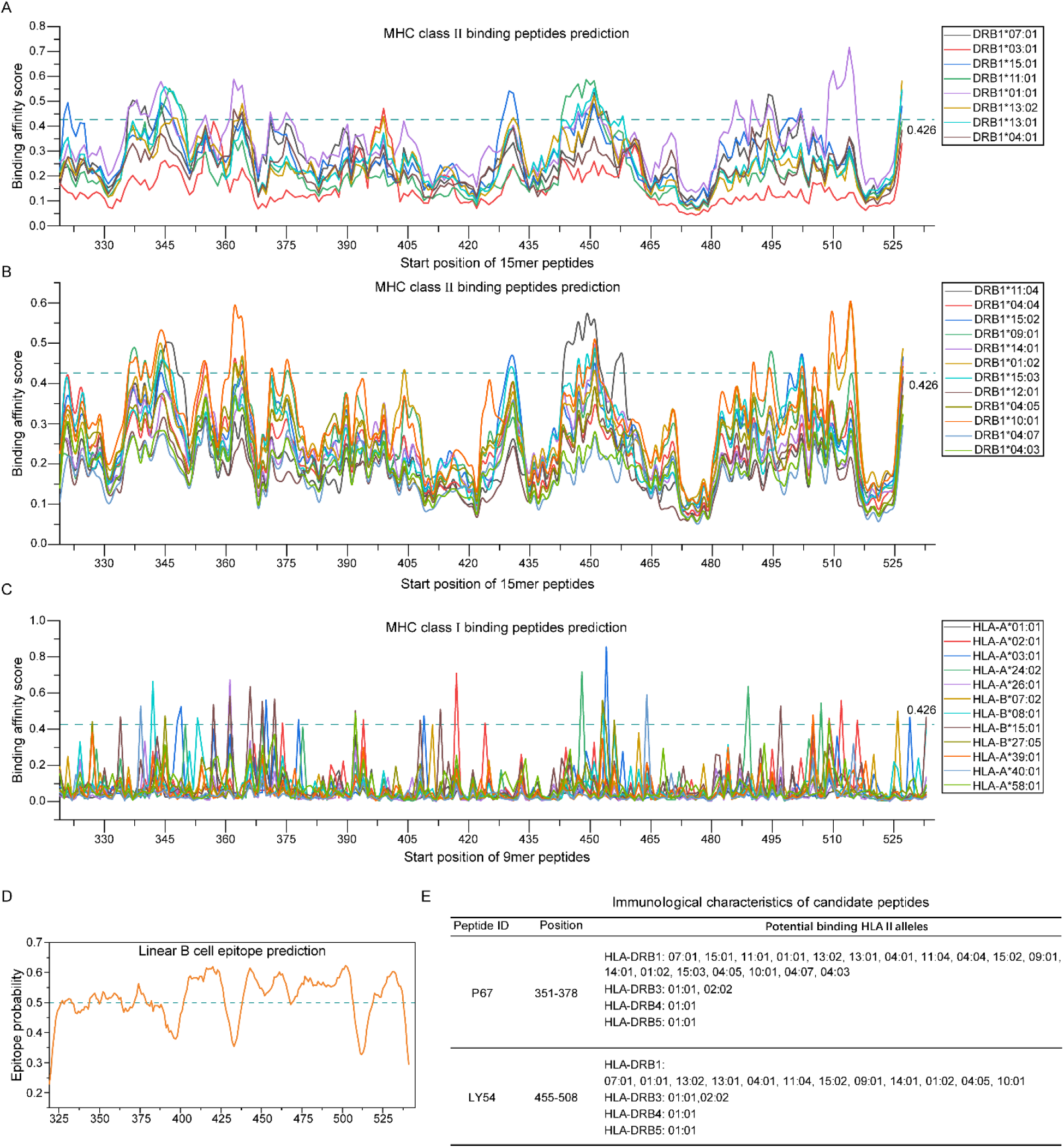
Immunogenicity prediction and candidate peptide selection for RBD. (**A** and **B**) The HLA class II binding peptides of RBD for major HLA class II types were predicted by NetMHCIIpan-4.0. Graphs were created with Origin 2019 software. (**C**) The HLA class I binding peptides of RBD for major HLA class I types were predicted by NetMHCpan-4.1. Graph was created with Origin 2019 software. (**D**) B-cell linear epitopes of RBD were predicted by BepiPred-2.0. Graph was created with Origin 2019 software. (**E**) The table shows that the potential binding HLA II alleles for LY54 and P67.

**Fig. S2.**
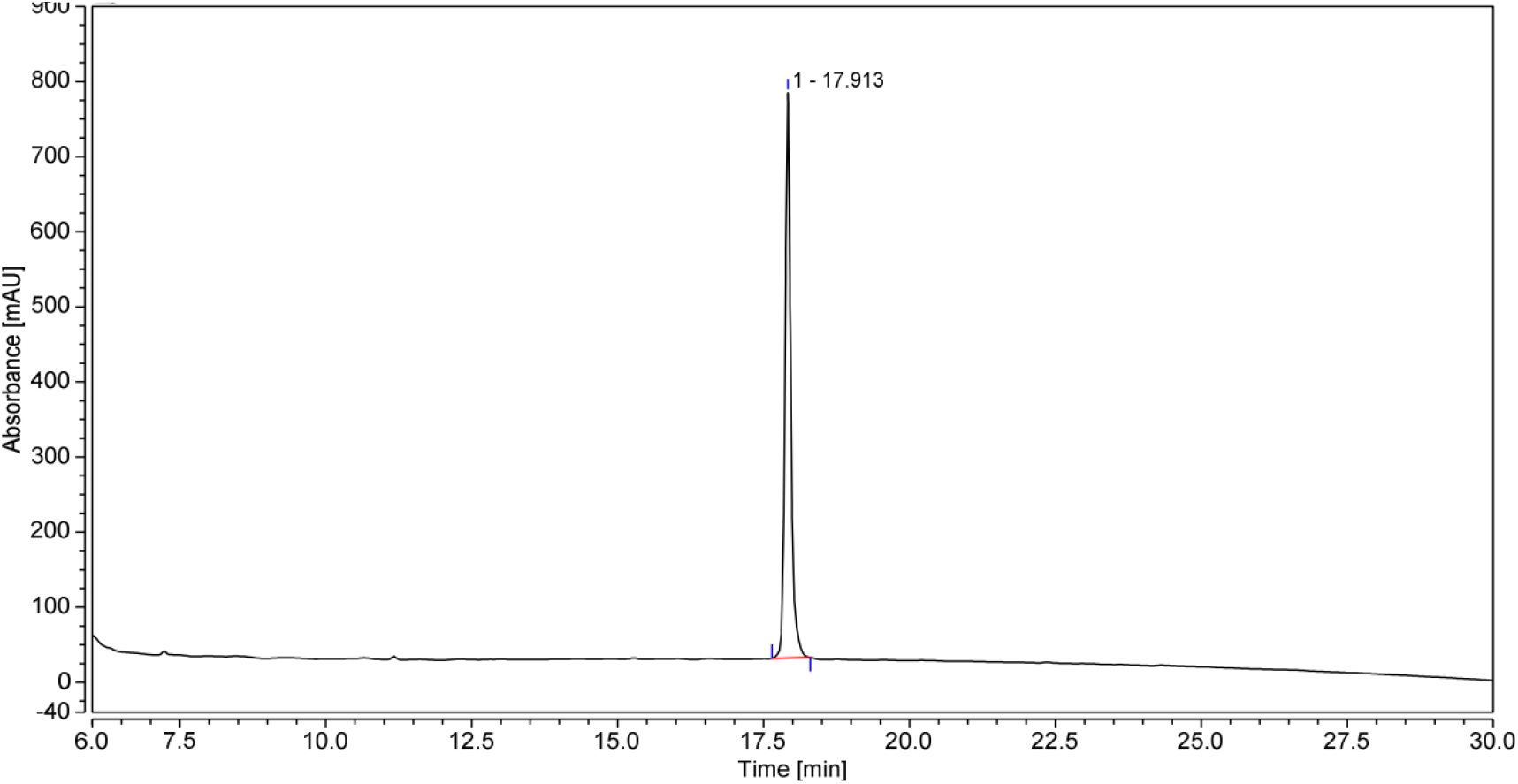
HPLC profile of purified LY54-101. The LY54-101 was purified by preparative C18 column and was obtained as a white powder after lyophilization. Purified LY54-101 was identified by HPLC.

**Fig. S3.**
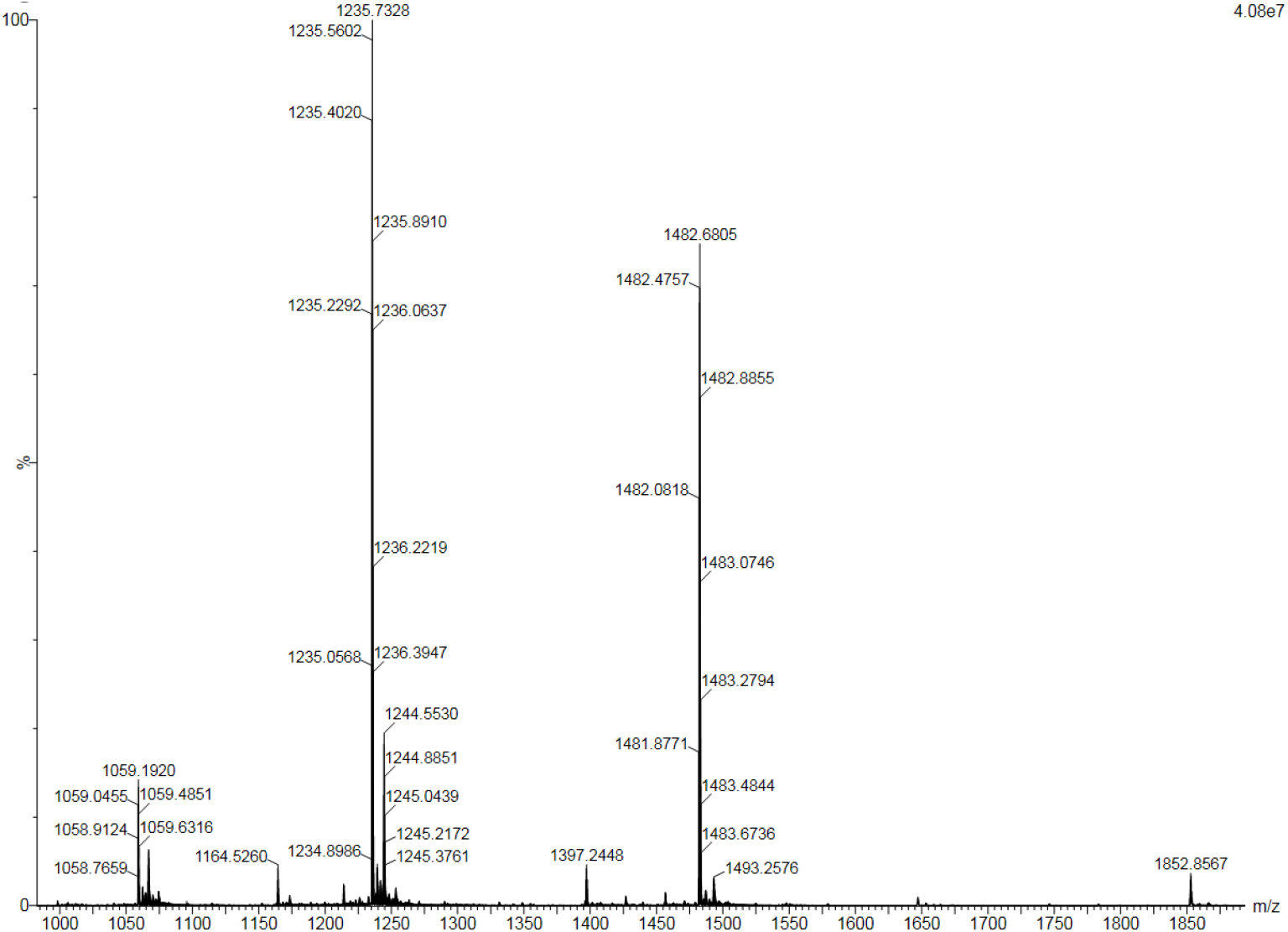
HRMS profile of purified LY54-101. The LY54-101 was purified by preparative C18 column and was obtained as a white powder after lyophilization. Purified LY54-101 was identified by HRMS.

**Fig. S4.**
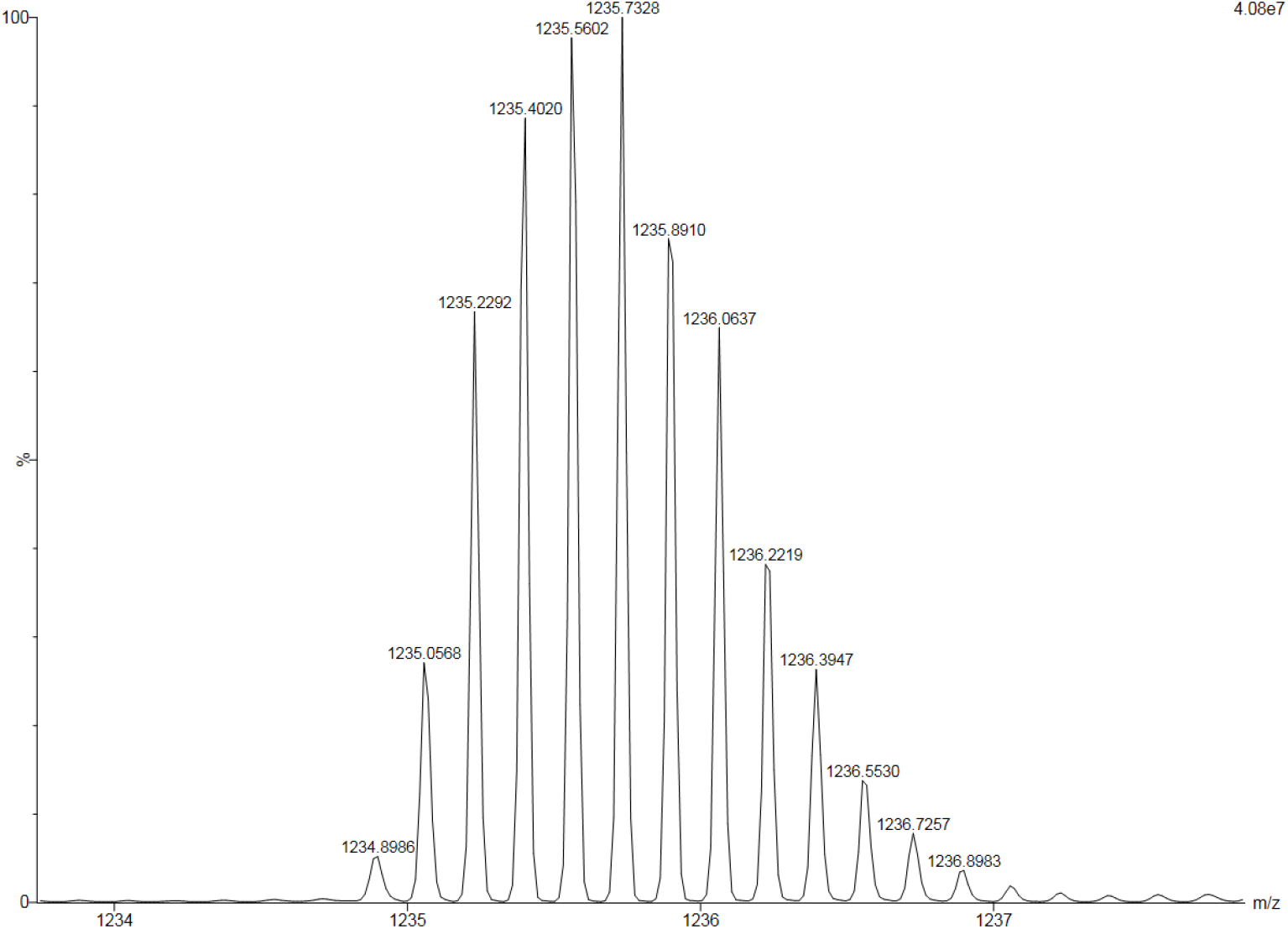
HRMS profile of purified LY54-101 (magnify [M+6H]6+ peak). The LY54-101 was purified by preparative C18 column and was obtained as a white powder after lyophilization. Purified LY54-101 was identified by HRMS.

**Fig. S5.**
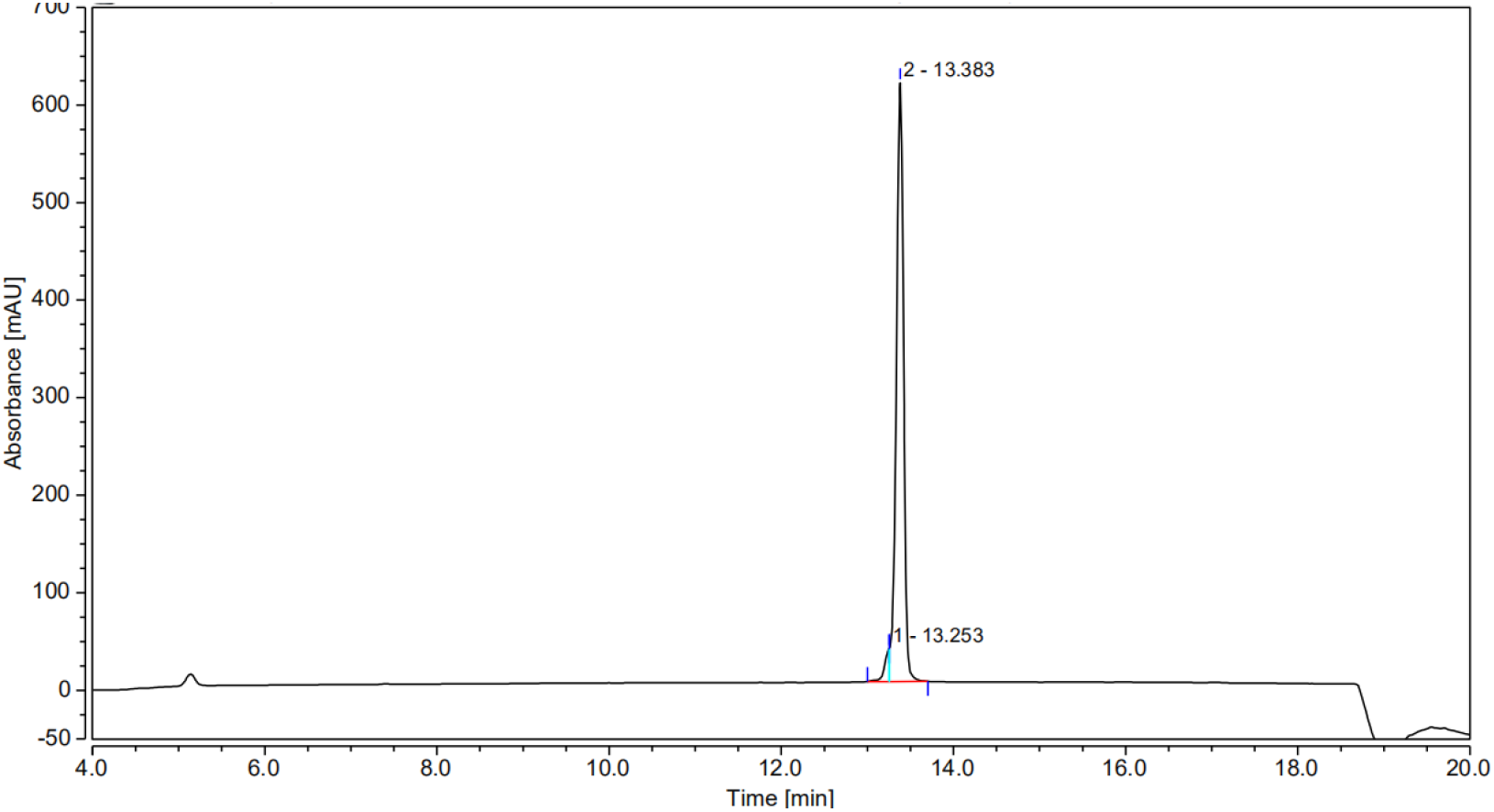
HPLC profile of purified P67-101. The P67-101 was purified by preparative C18 column and was obtained as a white powder after lyophilization. Purified LY54-101 was identified by HPLC.

**Fig. S6.**
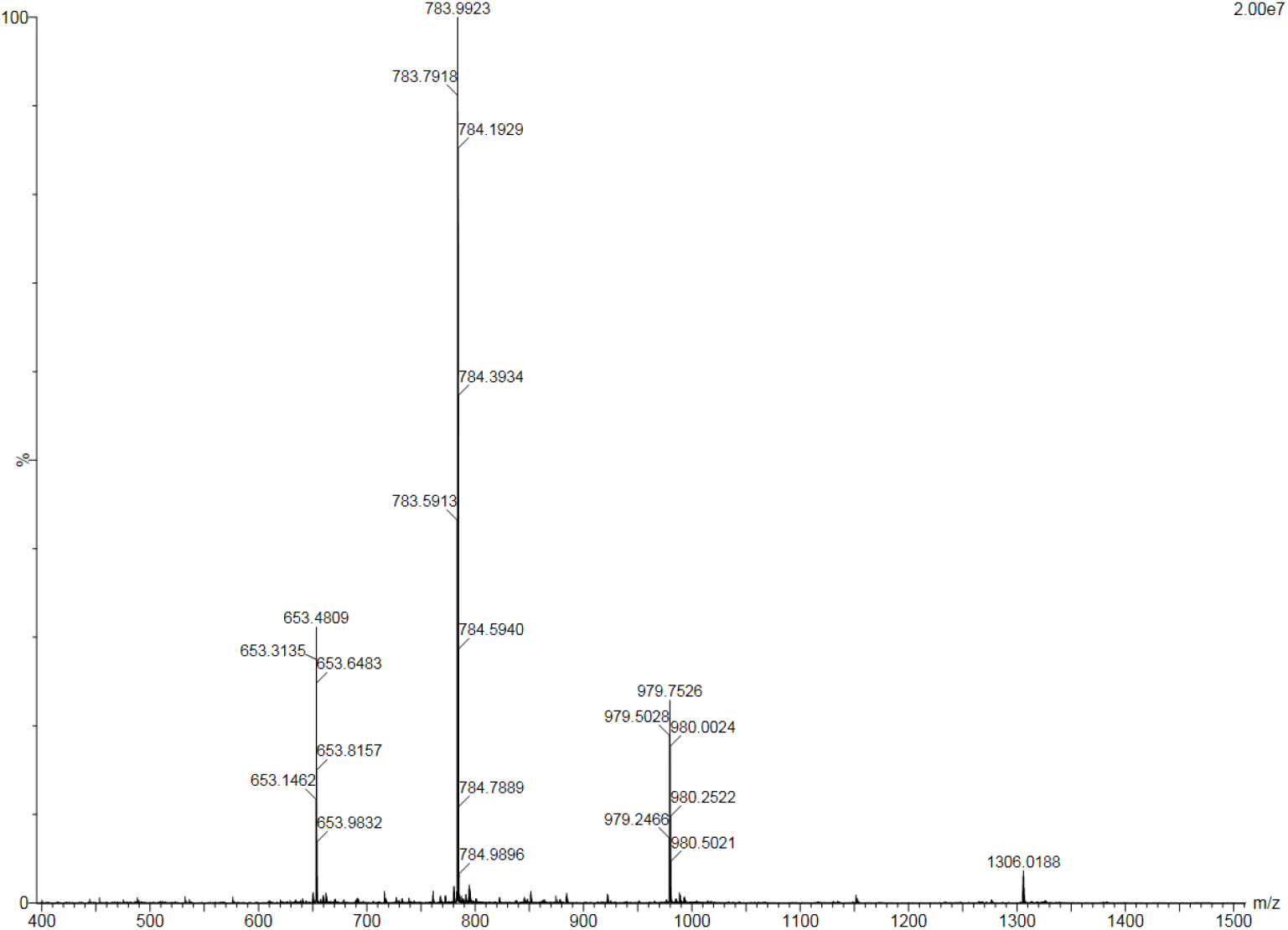
HRMS profile of purified P67-101. The P67-101 was purified by preparative C18 column and was obtained as a white powder after lyophilization. Purified LY54-101 was identified by HRMS.

**Fig. S7.**
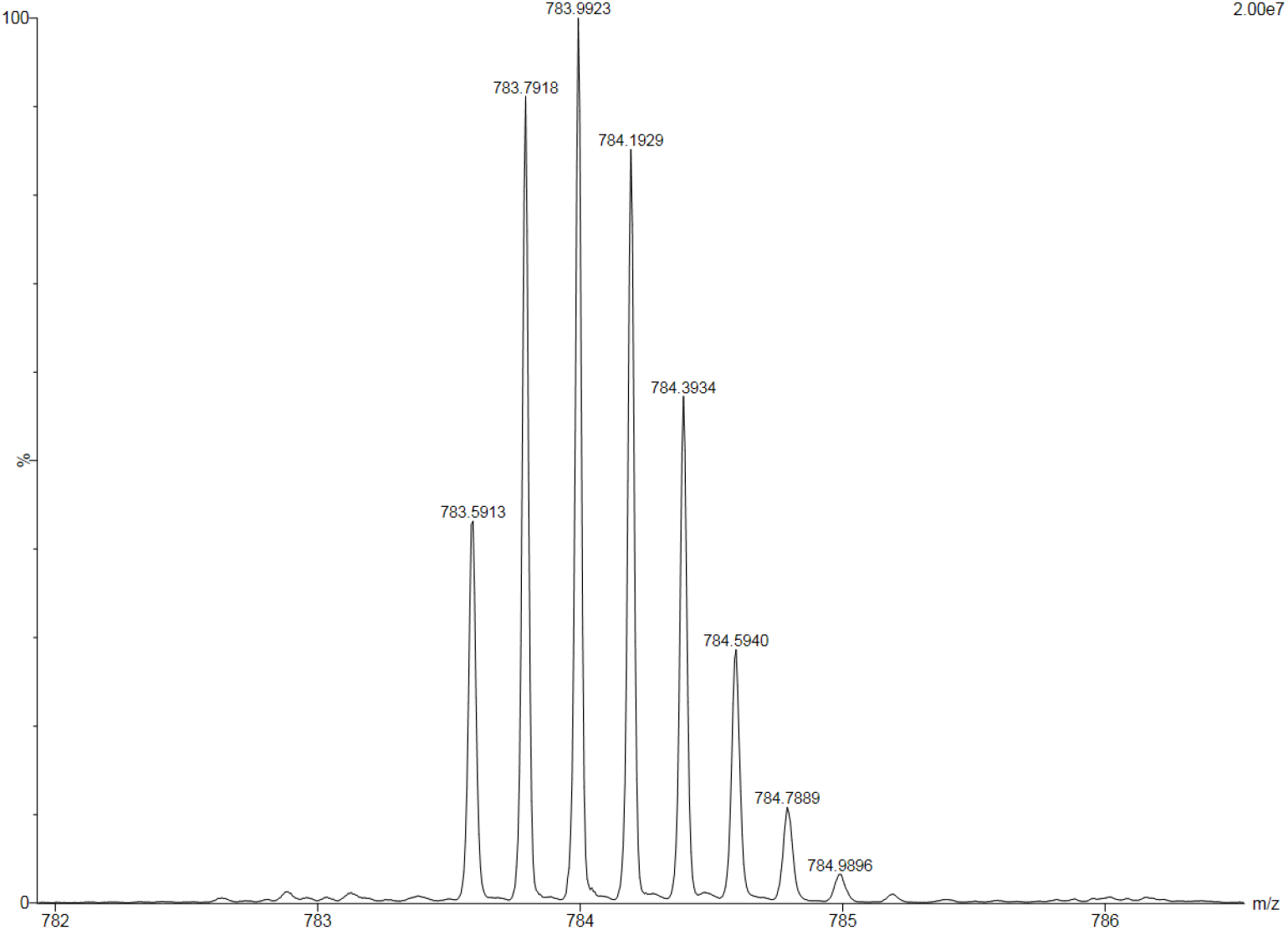
HRMS profile of purified P67-101 (magnify [M+5H]5+ peak). The P67-101 was purified by preparative C18 column and was obtained as a white powder after lyophilization. Purified LY54-101 was identified by HRMS.

**Fig. S8.**
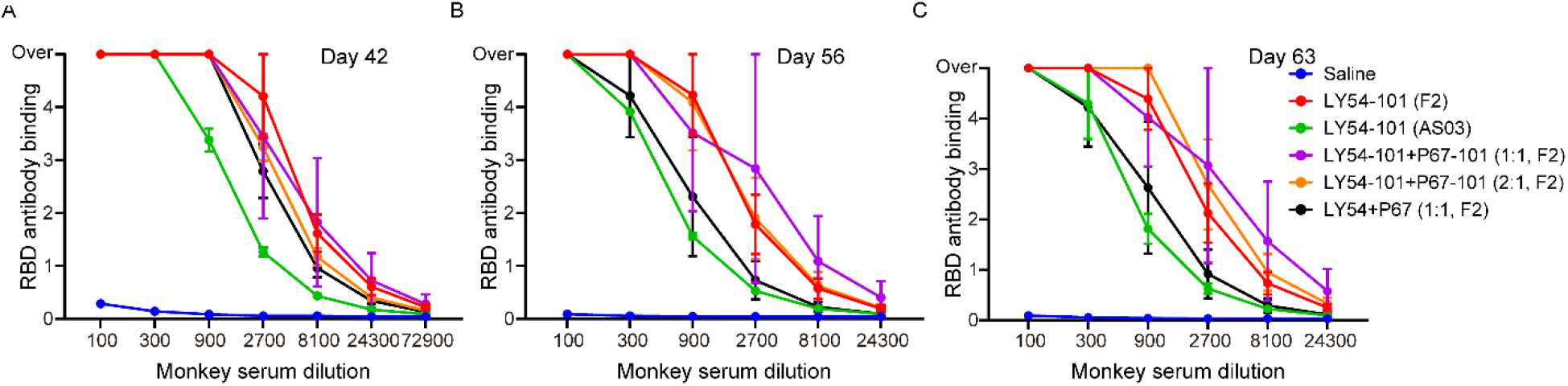
RBD binding antibodies in vaccinated cynomolgus monkeys. (**A** to **C**) The cynomolgus monkeys (n = 2) were immunized with peptide vaccines in the following groups: saline, LY54-101 (F2), LY54-101 (AS03), LY54-101+P67-101 (1:1, F2), LY54-101+P67-101 (2:1, F2) and LY54+P67 (1:1, F2) for three doses at days 0, 14 and 28. Sera were collected from the monkeys 42 (A), 56 (B) and 63 (C) days after the first dose of vaccine and the levels of RBD-specific antibody were tested for different serum dilutions using ELISA. The data are presented as the mean ± SEM.

